# Enhanced super-resolution microscopy by extreme value based emitter recovery

**DOI:** 10.1101/295261

**Authors:** Hongqiang Ma, Wei Jiang, Jianquan Xu, Yang Liu

**Author notes:** &.

## Abstract

Super-resolution localization microscopy allows visualization of biological structure at nanoscale resolution. However, the presence of heterogeneous background can degrade the nanoscale resolution by tens of nanometers and introduce significant image artifacts. Here we develop a new approach, referred to as extreme value based emitter recovery (EVER), to accurately recover the distorted fluorescent emitters from heterogeneous background. Through numerical simulation and biological experiments, we demonstrate that EVER significantly improves the accuracy and fidelity of the reconstructed super-resolution image for a wide variety of imaging characteristics. EVER requires no manual adjustment of parameters and is implemented as an easy-to-use ImageJ plugin that can immediately enhance the quality of super-resolution images. Our method paves the way for accurate nanoscale imaging of samples with heterogeneous background fluorescence, such as thicker tissue and cells.

## INTRODUCTION

Significant advance in localization-based super-resolution imaging techniques (also known as STORM, PALM or fPALM) ^1–4^ have revolutionized the field of light microscopy and allows visualization of the previously invisible molecular structures at a resolution of ~20 nm. Their super-resolved imaging capability is achieved by precise localization of individual fluorescent emitters with nanometer accuracy. However, in many biological experiments, the fluorescent emitters can be distorted or obscured by the heterogeneous noisy background, which can introduce inaccuracies up to tens of nanometers. The inaccuracy can significantly compromise the resolution and quality of the reconstructed super-resolution image in forms of image artifact (i.e., misrepresentation of samples’ structure) and localization bias (i.e., shifting the true positions of localized emitters) ^5–8^. Such limitation restricts the application of super-resolution localization microscopy to those thin cells and tissue where the background is relatively uniform. To achieve state-of-the-art super-resolution imaging ^5,6,8^ in a wide variety of biological samples (e.g., thicker tissue and cells), accurate emitter recovery from the heterogeneous background fluorescence is critical.

Earlier attempts to recover fluorescent emitters rely on conventional image processing methods, such as spatial filtering (e.g., rolling ball filter ^5,9^) and temporal filtering (e.g., temporal median filter ^10,11^). These approaches are not based on rigorous estimation models for super-resolution localization microscopy, and all have serious limitations in practice. Spatial filters lack robustness to background structure and emitter size. Temporal median filter suffers from serious over-estimation that can suppress the emitter intensity and size, which can severely reduce the emitter recall rate and affect the localization accuracy, especially in axial dimension^11^.

Here, we investigated the features of the composite signal in super-resolution localization microscopy and developed EVER, an accurate emitter recovery method using an extreme value-based estimation model. Our method accurately separates the fast varying emitter signals from the slow varying background signal without restrictions present in conventional methods, and can significantly reduce image artifacts and improve the accuracy and fidelity of super-resolution localization microscopy. We demonstrate the superior performance of EVER over conventional methods for a wide range of imaging characteristics using dataset from numerical simulation and biological experiments with tissue and cell samples. Moreover, we have implemented EVER as an easy-to-use ImageJ plugin to help the users to immediately improve the performance of their super-resolution localization microscopy.

## RESULTS

### 1. Extreme Value based Emitter Recovery (EVER)

Super-resolution localization microscopy requires sequential imaging of a subset of the densely labeled fluorophores at each image frame. During this process, heterogeneous background fluorescence may severely distort the detected emitters (Fig. 1a). To accurately recover the distorted emitters, we propose a new model, EVER, which uses pixelwise extreme value of an image sequence to separate the emitters from the background signal. The main procedures of EVER can be divided into two steps: (1) segment the raw image stack to a series of image sub-stack along the temporal axis and calculate the pixelwise minimum value for each image sub-stack (Fig. 1b); (2) estimate the background from the temporal minima based on our derived relationship function to recover the emitters (Fig. 1c).

The underlying principle of EVER to accurately recover emitters from background comes from the extreme value statistics. In super-resolution localization microscopy, the acquired composite signal composed of the fast-changing emitters and the slowly varying background, can be modeled as a Poisson distribution. In an extreme case with ultra-sparse emitter signal, the composite signal is nearly equivalent to the background signal, which can be well estimated by temporal median or mean value (illustrated by the green dashed line in Fig. 1b) and easily separated from the emitter signal. But in practice, a significantly higher emitter density is often employed for a satisfying imaging speed. For example, when the probability of emitter occurrence is 50%, the temporal median or mean value is significantly skewed towards the composite signals (illustrated by the green solid line in Fig. 1b), indicating its tendency to seriously over-estimate the background. In contrast, the extreme value 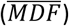 remains relatively stable regardless of the probability of emitter occurrence (illustrated by the red dashed and solid lines in Fig. 1b), suggesting its inherent robustness. However, the temporal minimum value is not equivalent to the actual background signal. We, therefore, derive an algebraic function (Fig. 1c) to establish the relationship to estimate the background signal directly from temporal minimum value and separate emitters from background (Figs. 1(d-f)).

**Fig. 1.**
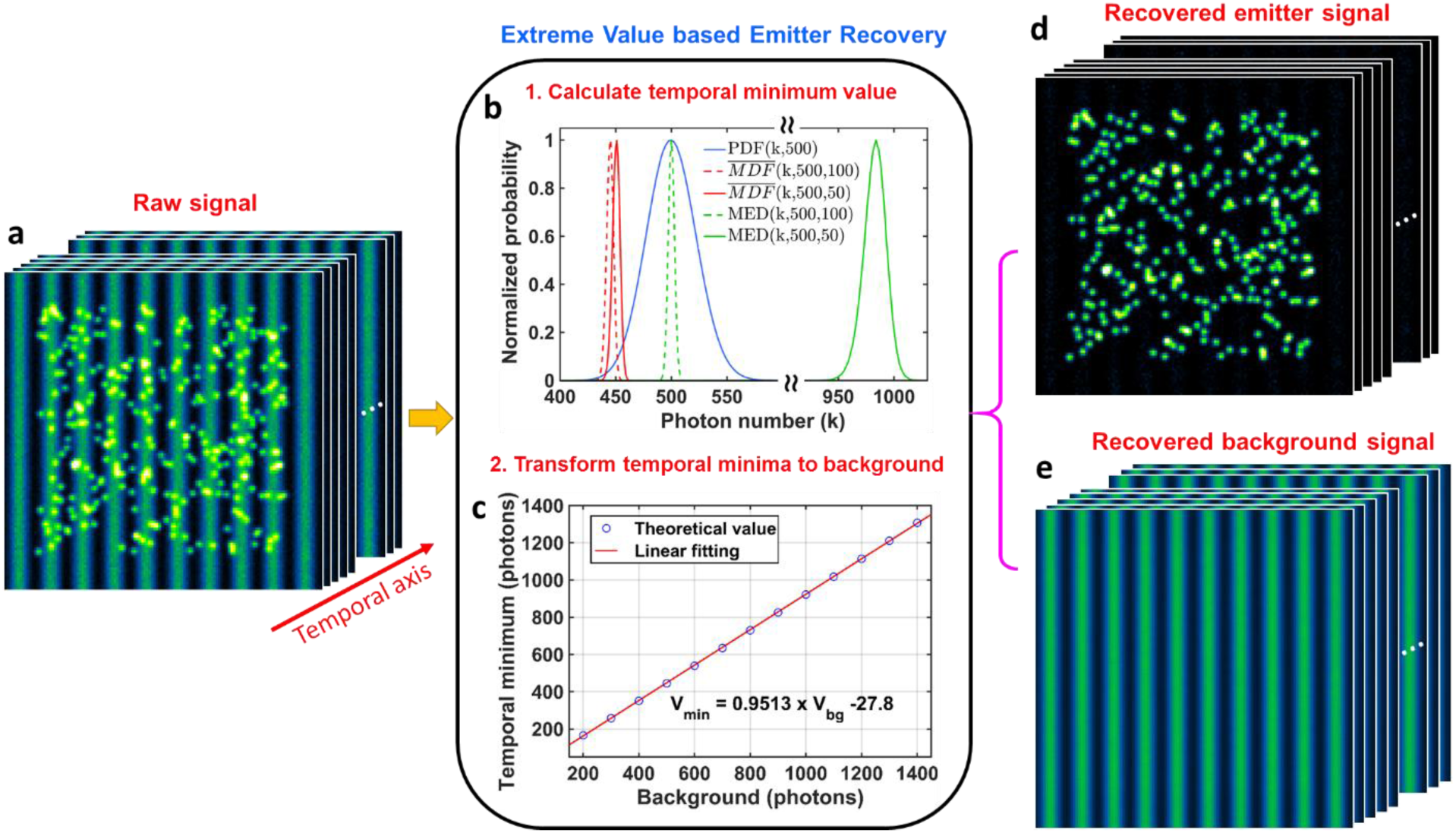
**(a)** An image sequence (stack) of raw images composed of emitters mixed with heterogeneous background. **(b-c)** The workflow of extreme value based emitter recovery (EVER), including **(b)** the calculation of temporal minimum value from each pixel and **(c)** transformation of temporal minimum value to the actual background value based on our derived algebraic function. **(d)** The recovered emitters and **(e)** background images. Note that, the blue line in (b) shows the probability density function (PDF) for a Poisson distribution with an expected background value of 500 photons. When probabilities of emitter occurrence is 50%, the mean of the probability distribution of temporal minimum value 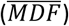 (red solid lines) only deviates by ~8 photons from the case with ultra-sparse emitters (red dashed line). While the temporal median value (MED) increases by ~480 photons for the same scenarios (green solid and dashed lines).

### 2. Validation of EVER against the ground truth of simulated dataset

We first validate the accuracy of EVER against the ground truth using simulated dataset that contains a rather complex imaging characteristics composed of heterogeneous background, emitter density, size and intensity. The detailed simulation parameters are described in Supplementary Methods. For each simulated dataset, the performance of our method is benchmarked against the ground truth and two conventional methods – temporal median filter (MED)^11^ and spatial rolling ball filter (RB)^9^. As shown in Fig. 2, our method accurately separates emitters from the noisy background, in which the recovered emitters show the best match with the ground-truth image. In comparison, the recovered emitters by MED exhibit apparent reduction in the intensity and size, due to the over-estimated background; the recovered emitters by the spatial filter of RB exhibit erroneous structures as the artifacts introduced by the heterogeneous background. We further quantify their image similarity of the recovered emitters using our method (EVER), MED and RB, as shown in Fig. 2c. Our method indeed best recovers the emitters with ~98% similarity ^12^ compared to the ground truth, while other methods only show <60% similarity with the ground truth. The similar performance holds for other imaging scenarios with various background structures, emitter density, size and intensity, as shown in Supplementary Figs. S1-S5.

**Fig. 2.**
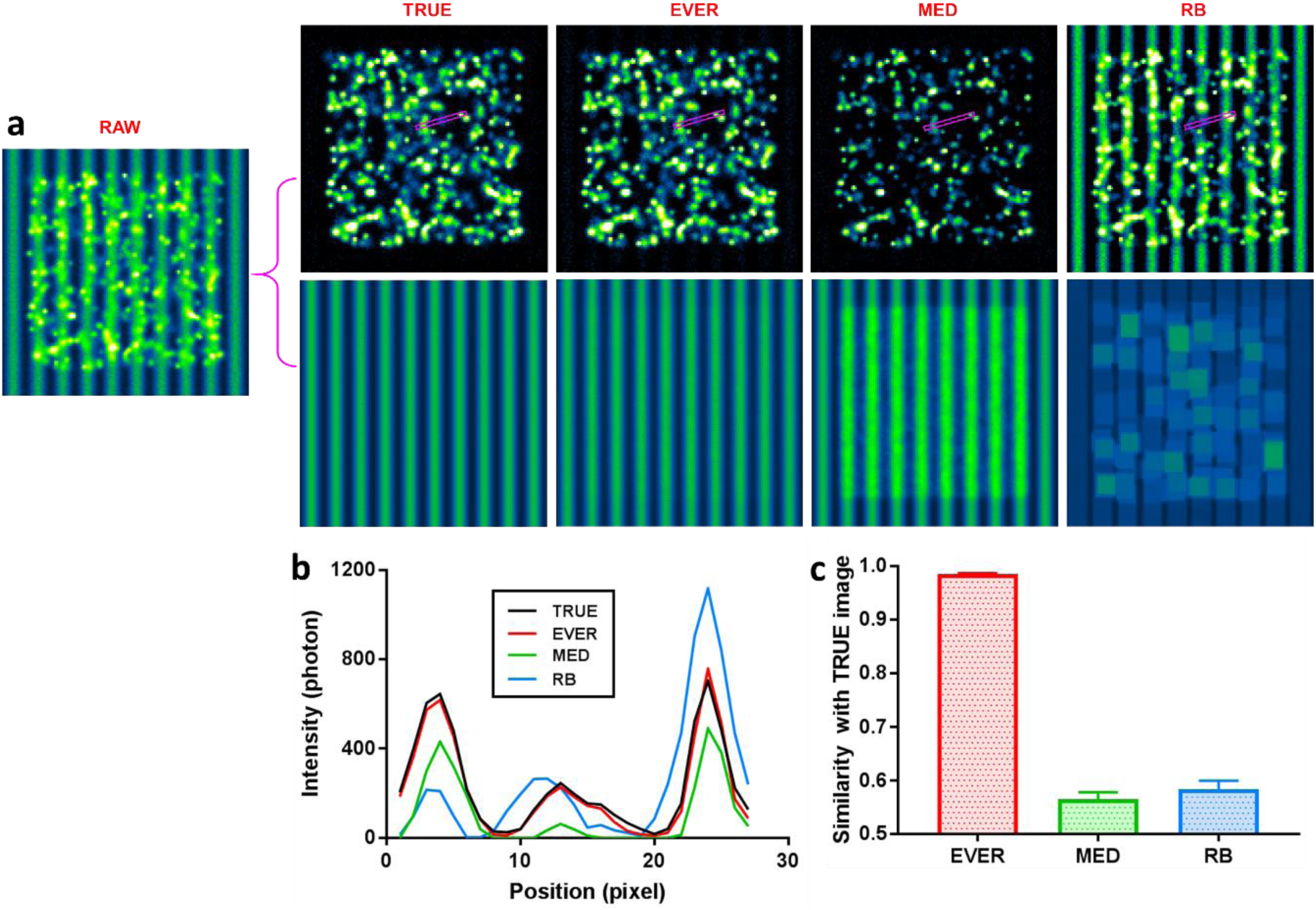
**(a)** A simulated raw image where the emitters are mixed with heterogeneous background, and the emitter and background images are recovered by ground-truth (TRUE), extreme value based emitter recovery (EVER), temporal median filter (MED) and spatial rolling ball filter (RB). **(b)** The intensity profile of the region in the magenta rectangular box of (a). **(c)** Image similarity between the recovered emitter images using EVER, MED and RB and the TRUE image.

### 3. Performance of EVER on the reconstructed super-resolution image

The accuracy of emitter recovery directly affects the quality of subsequent super-resolution image reconstruction. As shown in Fig. 3, we compared the performance of EVER against the conventional methods of temporal median filtering (MED) and spatial rolling ball filtering (RB) based on the simulated datasets with the known ground truth. Figure 3(a) shows the simulated single-frame raw image (a1) with three emitters on top of a heterogeneous background, and the recovered emitter images from the ground truth (a2) and using EVER (a3), MED (a4) and RB (a5), respectively. The recovered emitters using EVER (a3) best resemble those from the ground truth (a2). While the emitter images recovered by MED (a4) show much smaller emitter size and lower emitter intensity, and those recovered RB (a5) show the residual background features. This observation is further confirmed in the corresponding cross-sectional profile shown in Fig. 3(c). The red solid line from EVER best tracks the profile from ground truth (black solid line), but the profile from the MED-recovered image (green) shows the over-estimated background with reduced emitter intensity and size, and the profile from the RB-recovered image (blue) shows the under-estimated background such as the region indicated by the black arrow.

Following emitter recovery, we reconstructed the super-resolution images reconstructed by ThunderSTORM^13,14^, rendered as the 2D image (Figs. 3(b, d-e)) and the 3D scattered plot of localized emitter positions (Fig. 3(d)). Without any processing, the heterogeneous background results in many erroneously localized emitters, shown as image artifacts in Figs. 3(b1, d1). After EVER, the reconstructed super-resolution image (Fig. 3(b3)) shows the closest match to the super-resolution image reconstructed from the ground truth (Fig. 3(b2)). Whereas other methods (MED and RB) suffer from apparent reduction in localization accuracy and image resolution. Table 1 compares the performance of emitter recovery on localization accuracy of the emitter position and size, quantified by localization bias and precision. Our method shows the best localization accuracy, comparable to that from the ground truth.

**Fig. 3.**
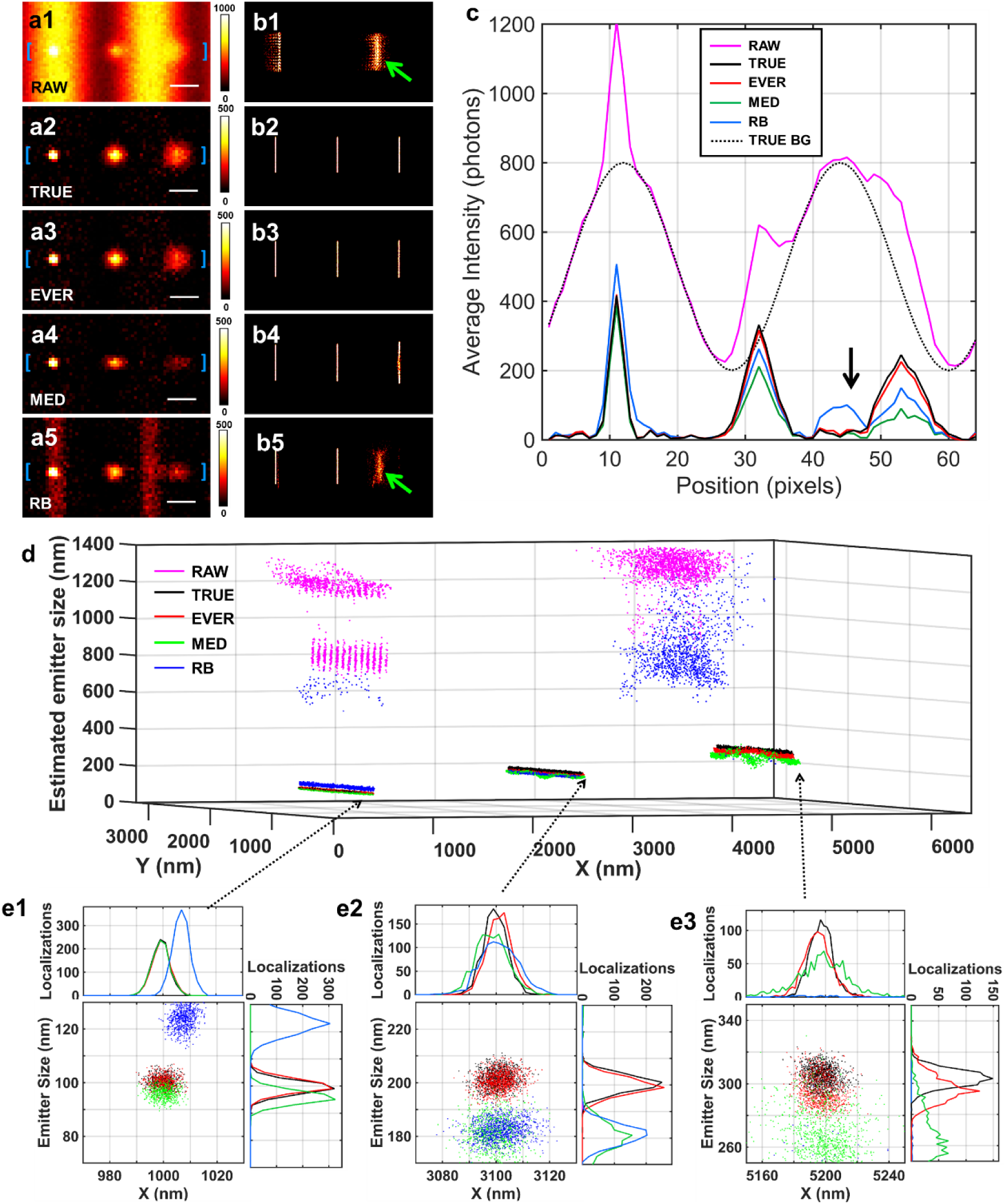
The comparison of different emitter recovery approaches using simulated dataset with various emitter size, emitter intensity and heterogeneous background. **(a)** The raw image (a1), the recovered emitters from the ground truth (TRUE) (a2), and using EVER (a3), MED (a4) and RB (a5). **(b)** The corresponding reconstructed super-resolution images using ThunderSTORM without additional processing (b1) and after emitter recovery using the ground truth (b2), EVER (b3), MED (b4) and RB (b5). Green arrows indicate the image artifact. **(c)** The corresponding intensity profiles (solid lines) from the regions between the blue bracket in (a1-a5) and the black dashed line (TRUE-BG) represents the profile from the ground-truth background. The black arrow indicates the erroneously recovered emitter signals by RB coming from the residual background. **(d)** The scatter plot of the localized emitters from the raw image without any processing (magenta) and with emitter recovery by the ground truth (black), EVER (red), MED (green) and RB (blue). Emitter localization was performed by ThunderSTORM. **(e1-e3)** The zoomed distribution of localized emitters in (c).

**Table 1.** Localization bias and precision* of the emitter positions (x-dimension) and size for recovered emitters from the ground truth (TRUE), EVER, MED and RB.

|  |  | Bias (nm) |  |  | Precision (nm) |  |  |
| --- | --- | --- | --- | --- | --- | --- | --- |
|  |  | Left emitter | Middle emitter | Right emitter | Left emitter | Middle emitter | Right emitter |
| **Lateral position** | TRUE | 0.4 | 1.5 | 1.9 | 2.7 | 3.4 | 5.5 |
|  | EVER | 0.9 | 2.1 | 3.3 | 2.8 | 3.6 | 7.1 |
|  | MED | 1.9 | 2.3 | 3.7 | 2.8 | 5.4 | 12.3 |
|  | RB | 9.1 | 2.2 | >50 | 4.4 | 4.2 | >50 |
| **Emitter size** | TRUE | 0.1 | 0.8 | 3.1 | 3.5 | 4.4 | 6.9 |
|  | EVER | 0.1 | 0.9 | 4.9 | 3.6 | 5.1 | 7.6 |
|  | MED | 5.0 | 21.3 | 40.6 | 3.7 | 6.3 | 12.8 |
|  | RB | 22.9 | 17.6 | >50 | 3.5 | 7.1 | >50 |

\*Note: The localization bias is defined as the mean value of the localization error, and the localization precision is defined as the standard deviation of the localization error.

### 4. Experimental results in tissue imaging

To demonstrate the performance of EVER on super-resolution imaging of biological samples, we first used an experimental dataset of imaging heterochromatin from tissue section. Super-resolution imaging of tissue section is especially challenging due to the presence of heterogeneous strong background mainly from tissue autofluorescence and stronger scattering; the imaging target of heterochromatin further complicates this problem due to the high emitter density from the densely packed heterochromatin. Figure 4(a1) shows the representative single-frame raw image that exhibits a strong heterogeneous background mixed with emitters. Figures 4(a2-a4) show the recovered emitters and the estimated background by EVER, MED and RB, respectively. Figures 4(c1-c4) show the reconstructed super-resolution images using emitters recovered by EVER, MED and RB, respectively. Evidently, the MED recovers less number of emitters with reduced intensity due to the over-estimated background (Fig. 4(a3)), which results in much lower recalled rate in the reconstructed super-resolution image (~1.9×10^5^ localized emitters) compared to other methods (>2.8×10^5^). On the other hand, the RB under-estimates the background and recovers erroneous emitters that introduce image artifacts that are not visible in the wide-field image (Fig. 4b), as indicated in the blue regions in Figs. 4(b, c4). In comparison, the reconstructed super-resolution image after our EVER (Fig. 4(c2)) indeed shows the best match with the overall structural features in the wide-field image without any image artifacts or loss of recovered emitters.

**Fig. 4.**
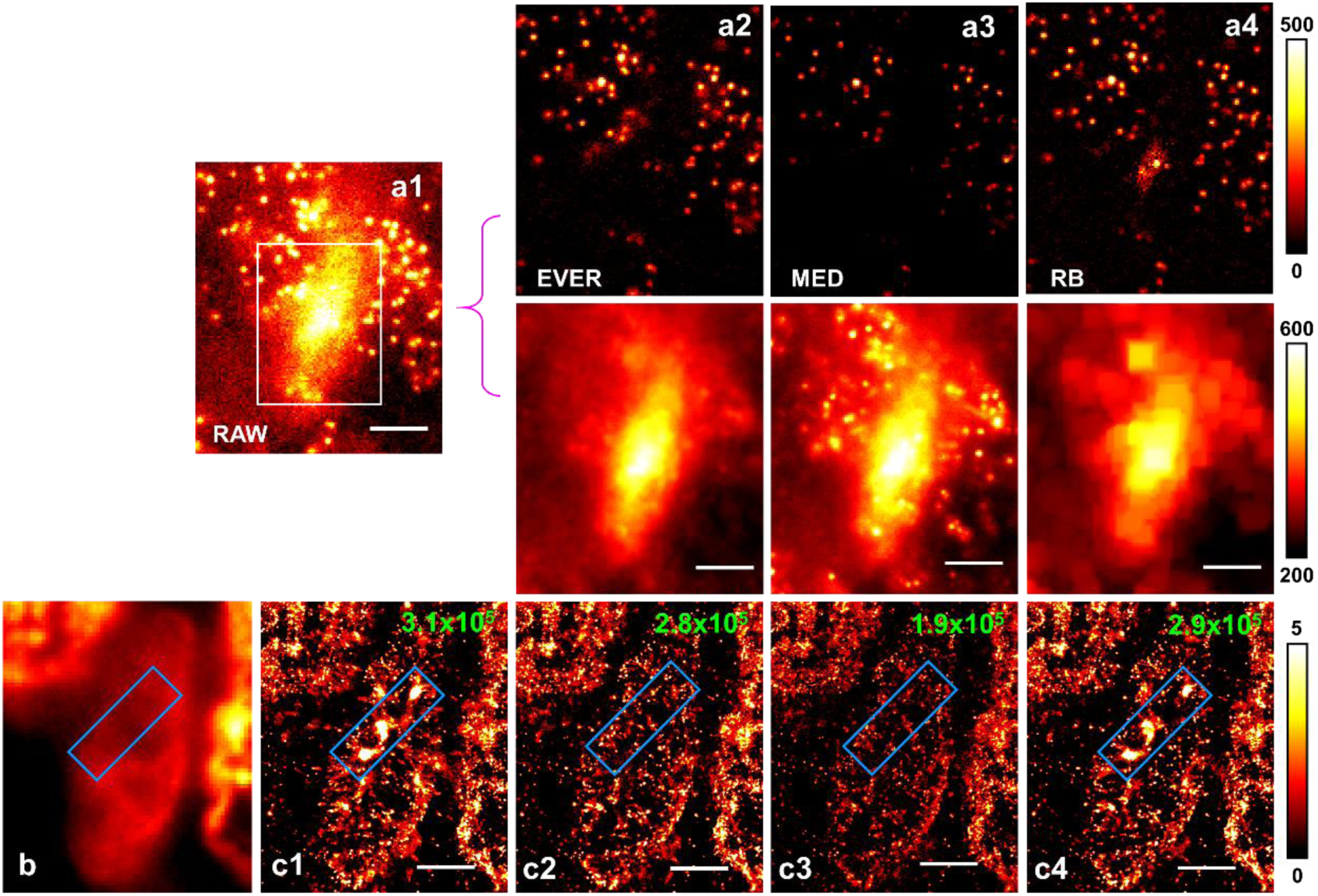
Performance of our extreme value-based emitter recovery (EVER) for super-resolution localization microscopy. (a1) A single-frame raw image of the heterochromatin (labeled by Alexa 647-conjugated antibody against H3K27me3) from a tissue section in the imaging condition for super-resolution localization microscopy and the recovered emitters and background using EVER (a2), MED (a3), and RB (a4). Scale bar: 4 µm. (b) A wide-field image and the super-resolution image of heterochromatin reconstructed by ThunderSTORM without any additional processing (c1) and with emitter recovery by EVER (c2), MED (c3), and RB (c4). Blue box indicate the image artifact caused by the heterogeneous background. The number of recalled emitters is shown on the upper right corner of each reconstructed image. Scale bar: 2 µm.

### 5. Experimental results in cell imaging

Although strong heterogeneous background is less common on imaging thin cells, it can be significant in multi-color dSTORM imaging when strong crosstalk (light from one color leaked into another color channel) is present. In our experiment, we labelled the mitochondria with Alexa647 and microtubule with Cy3B and a four-band dichroic mirror and emission filter were used in our microscopy setup. Figure 5(b1) shows strong non-uniform background due to the cross-talk from the Cy3B-labeled microtubule when imaging the mitochondria. Figures 5(b2-b4) show the recovered emitters by EVER, MED and RB. Similar to our previous findings, the MED-recovered image (Fig. 5(b3)) shows significantly less number of emitters and lower emitter intensity due to background over-estimation, while RB-recovered image (Fig. 5(b4)) exhibits the residual background features as a result of background under-estimation. In comparison, our method (Fig. 5(b2)) recovers more emitters of various sizes (than MED), without apparent background features. In the subsequent reconstructed super-resolution images of mitochondria, we found that the reconstructed super-resolution images without additional processing (Figs. 5(c1, e1)) and with RB-based emitter recovery (Figs. 5(c4, c4)) exhibit apparent artifacts that are not seen in the wide-field image (as indicated by the blue box in Figs. 5(d)). Both EVER (Figs. 8(c2, e2)) and MED-based emitter recovery (Figs. 5(c3, e3)) match the structural features seen in the wide-field image (Figs. 5(d)), but the reconstructed image after MED-based emitter recovery shows a significantly lower number of recalled emitters. Although MED-based recovery eliminates the image artifacts caused by non-uniform background, but significantly sacrifices the emitter intensity, emitter size and recall rate due to the over-estimated background, consistent with our previous simulation and experimental results.

**Fig. 5.**
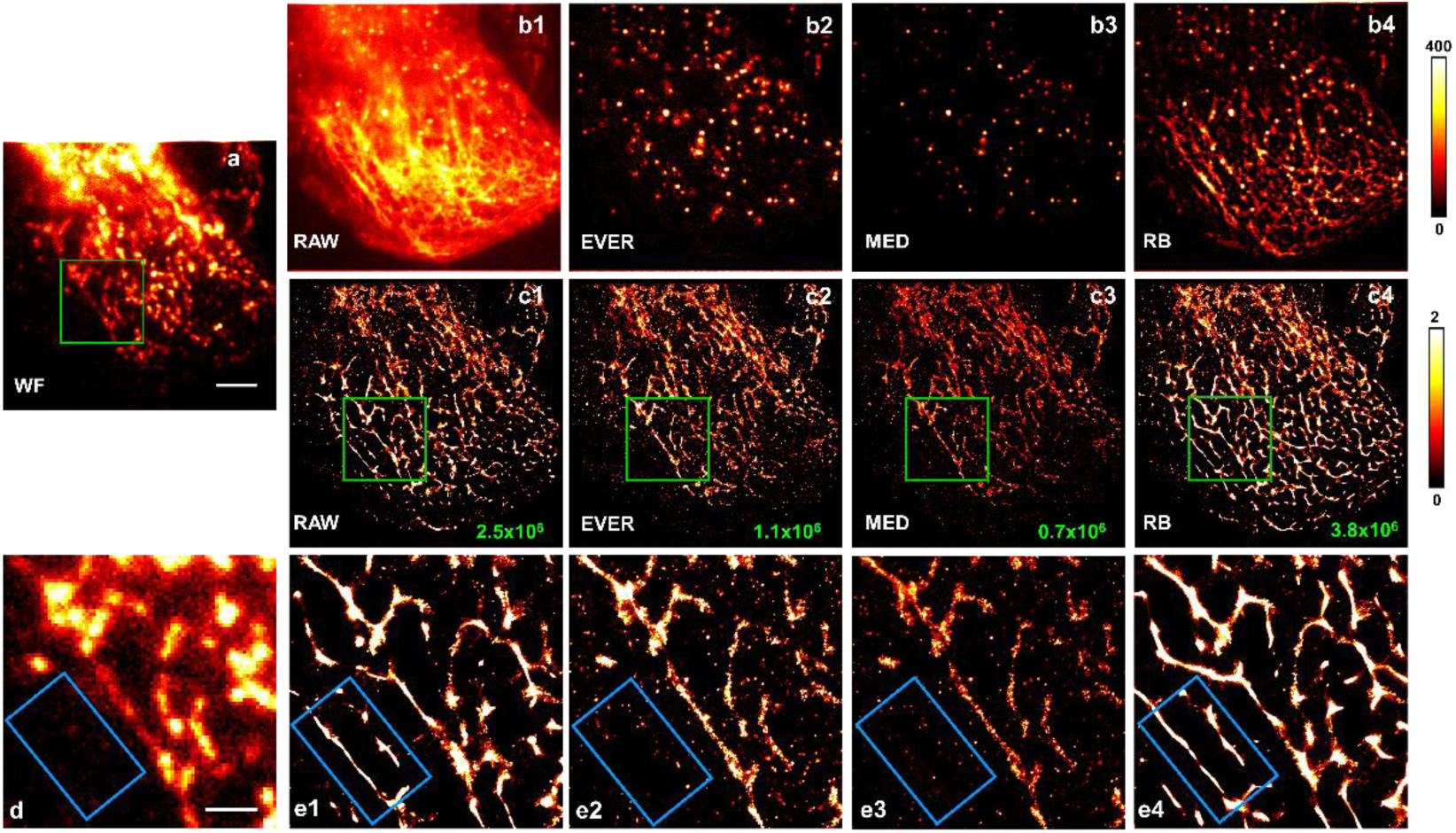
Comparison of different emitter recovery approaches for cell imaging using super-resolution localization microscopy. (a) The wide-field (WF) image of the experimental mitochondria dataset. (b1-b4) The single-frame raw image and the corresponding recovered emitters using EVER (b2), MED (b3), and RB (b4). Scale bar: 5 µm. (c2-c4) The reconstructed super-resolution image of mitochondria (labeled by Alexa 647-conjugated antibody) by ThunderSTORM without any additional processing (c1) and with emitter recovery by EVER (c2), MED (c3), and RB (c4). The number of recalled emitters is shown in the lower right corner of each reconstructed image. (d, c1-c5) The zoomed regions within the green boxes of the wide-field image (a) and reconstructed super-resolution images in (c1-c4). Scale bar: 2 µm.

### 6. Computation speed

We also compared the computation speed of different methods using ImageJ (Intel Core i7-4790 @ 3.6 GHz, only one thread was used) for our experiment dataset in Fig. 4 (image size: 128×152 pixels, frame number: 20000 frames). Our method (EVER) takes ~9 seconds, RB takes ~115 seconds, and MED takes ~53 second. Therefore, besides the superior performance demonstrated above, our method is also significantly faster that RB and MED by a factor of 12 and 5, respectively. This speed is sufficient for online analysis for the increasingly large dataset of super-resolution localization microscopy.

## CONCLUSION

In conclusion, we present an extreme value based emitter recovery method, EVER, that improves the fidelity and the resolution of super-resolution localization microscopy with a robust performance for a wide range of imaging characteristics. In our approach, we use a time-domain statistical model that is more suitable than conventional spatial filters to separate the fast-changing emitters from the slowly varying background^11^. However, it is rather counter-intuitive to use the extreme value for background estimation, as it is traditionally considered to be less robust compared to temporal median or mean filter in conventional imaging processing. We demonstrate that in super-resolution localization microscopy, it exhibits superior robustness to separate the fast-varying emitters from the slowly-varying background. We also establish, for the first time to our knowledge, a simple algebraic relationship to link the temporal minimum value to the actual background signals. We validate the accuracy and robust performance of our method against the ground truth using simulated datasets with a wide range of emitter and background characteristics. We also demonstrate that EVER enables robust and accurate nanoscale imaging in challenging scenarios for super-resolution localization microscopy, such as imaging densely packed structures in tissue slices with heterogeneous bright background and strong color cross-talk, where conventional methods result in image artifacts, compromised image resolution, and reduced emitter recall rate. We implement this method as an easy-to-use ImageJ plugin, which can be directly applied to the dataset from sparse, high-density, 3D super-resolution localization microscopy to immediately enhance their image resolution and reduce artifacts without any manual parameter adjustment.

## METHODS

### 1. Theoretical basis and implementation of extreme value-based emitter recovery (EVER)

In super-resolution localization microscopy, as the read noise, dark noise and the corrected fixed pattern noise can be neglected for most advanced cameras (e.g. sCMOS cameras ^15–17^), the acquired signal can be modeled as a Poisson distribution. The probability distribution function (PDF), cumulative distribution function (CDF) and minimum value distribution function (MDF) of a Poisson distribution can be described by the following equations:

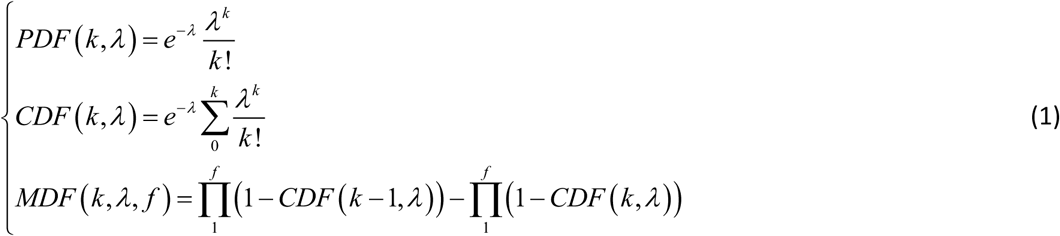

where k is the photon number, λ is the expected average photon number for each pixel and f is the length of signal used to estimate the minimum value.

The distribution of MDF can be further narrowed (reduced standard deviation), if a spatial mean filter (e.g., 3 × 3 mean filter) is applied. The mean of the temporal minimum value distribution function 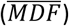 can be described by Equation 2:

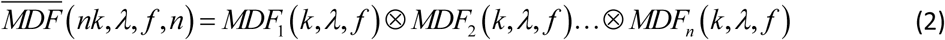

where n is the number of pixels being averaged.

Equation 2 is rather complicated and difficult to implement for large dataset from super-resolution localization microscopy. We thus derived a simple linear equation (Eq. 3) to relate the pixelwise temporal minima (V_MIN_) to the corresponding background (V_BG_). Based on Eq. 2, the expected theoretical temporal minimum values (100 frames) for a wide range of background levels (200-1400 photons) are shown in Fig. 1c. We obtain a perfect fit (R^2^>0.9999, residual error < 4 photons) for the background levels commonly seen in the localization-based super-resolution imaging dataset:

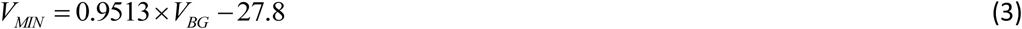

where *V*_*BG*_ is the true background and *V*_*MIN*_ is the temporal minimum value.

However, in some cases, the background fluorescence undergoes a slow variation (e.g., decay or rise) rather than remaining constant, even for a small subset of image frames (e.g., 100 frames). To further improve the robustness of our method, we modified Eq. 3 by incorporating the impact of different background variation rates, which is shown in the following Eq. 4 (as shown in Supplementary Fig. S6).

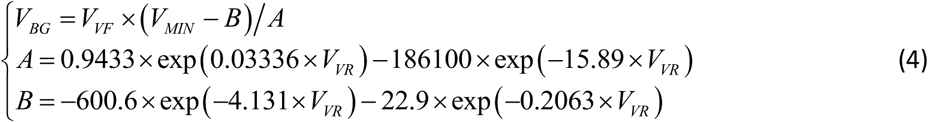

where V_f_ is the average intensity for the image set; V_VF_ is the variation factor for the f_th_ frame, defined as V_f_/min(V_1_:V_100_); V_VR_ is variation ratio, defined as the average value of the V_VF_. The variation of A and B along different variation ratios can be well approximated by a simple two-term exponential function, as shown in Figs. S6(b-c). When V_VR_ ≈ 1 (little variation), Eq. 4 can be reduced to Eq. 3.

### 2. Numerical simulation

To mimic a wide range of image characteristics in super-resolution localization microscopy, we simulated a series of image sets composed of spatially non-uniform background and emitters with various emitter density, size and intensity. The image size was set to be 128×128 pixels with a pixel size of 100 nm, and the emitters were randomly distributed in the central 100×100 pixels. For each image frame, the fluorescence signal is modeled as a distribution of emitters convolved with a point spread function and a spatially-varying background^18^:

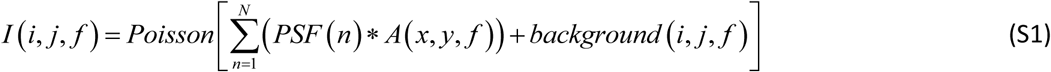

where f is the frame number, (i, j) is the coordinate of each pixel on the image, and A is the intensity of the nth emitter, (x, y) is the lateral position of the nth emitter, PSF(n) is the point spread function of the nth emitter and N is the total number of emitters in the image.

### 3. Image reconstruction

To compare our method with conventional methods for emitter recovery, the temporal median filter (MED) is implemented following the literature ^11^, and rolling ball filter (RB) is implemented by using ImageJ plugin (Process>> Subtract background) with a radius of 5 pixels. For super-resolution image reconstruction, ThunderSTORM ^13^ was used for all the simulation and experiments in this study. Wavelet filter was selected for image denoising and single-molecule least-squares Gaussian function fitting was used for localization. We acknowledge that our method improves the localization accuracy affected by non-uniform background, but it does not improve the localization precision affected by the increased Poisson noise from a high background. Furthermore, as we used least-square Gaussian localization algorithm for our super-resolution image reconstruction, we directly remove the background from our raw image set for the subsequent image reconstruction. However, if maximum likelihood estimator (MLE) is used, the estimated background image is needed for precise calculation.

### 3. Tissue sample preparation

#### A. Immunofluorescence staining of tissue section

A 3 µm-thick tissue section was cut from formalin-fixed, paraffin-embedded (FFPE) tissue block of a colon tissue and placed on a No. 1.5 coverslip. The tissue section was first deparaffinized in xylene and rehydrated in ethanol with graded concentration (100%, 95%, 70%, and 50%) and finally in distilled water. Next, heat-induced antigen retrieval was performed in the pre-heated Tris-EDTA buffer solution in microwave oven, then cooled down at room temperature. To block against non-specific binding, the section was incubated with a blocking solution containing 3% BSA and 0.2% Triton X-100 diluted in PBS for 1 hour at room temperature. The primary antibody (rabbit polyclonal H3K27me3, Cat. 07-449, Millipore) was diluted to be 1:300 in a solution containing 10mM glycine, 0.05% Tween 20, 0.1% Triton X-100, 0.1% hydrogen peroxide and 3% BSA in PBS and incubated at 4°C overnight, followed by Alexa Fluor 647 conjugated goat-anti-rabbit secondary antibody at room temperature for 2 hours and then washed with PBS.

#### B. Imaging buffer

The coverslip that containing the tissue section was glued to a plastic petri dish. The 70% of 2,2- thiodiethanol (TDE, Sigma-Aldrich) in PBS was used to optically clear the tissue section for at least 10 minutes before imaging. The imaging buffer was prepared fresh by mixing GLOX, 2-mercaptoethanol (βME, Sigma-Aldrich), Cyclooctatetraene (COT, Sigma-Aldrich) and TDE buffer B at a ratio of 1:1:1:97 before imaging. In brief, the GLOX was mixed with 200 µl Buffer A (0.5mL 1M Tris (pH = 8.0) + 0.146 g NaCl + 50 mL H_2_O), 14 mg Glucose Oxidase (Sigma-Aldrich), 50 µl Catalase (17 mg/mL catalase as prepared by dissolving 0.85 mg Catalase in 50 µl Buffer A, Sigma-Aldrich) and TDE Buffer B (2.5 mL 1M Tris (pH 8.0) + 0.029 g NaCl + 5 g Glucose + 17.5 mL H_2_O + 30 mL TDE).

#### C. Cell sample preparation

MEF cells were maintained in DMEM medium supplemented with 10% FBS. Cells were plated onto a glass-bottom dish (World Precision Instruments, FD3510) at an initial confluency of 50% and cultured overnight to let the cells attach. To perform immunostaining, the cells were first pre-extracted for 30 s in 0.5% Triton X-100 (Triton) in BRB80 (80 mM PIPES, 1 mM MgCl2, 1 mM EGTA, pH=6.8) supplemented with 4 mM EGTA, washed in PBS, fixed with cold Methanol for 10 minutes. The cells were then incubated with primary antibodies (rabbit anti-alpha tubulin antibody, abacm 18251; mouse anti COX IV Mitochondrial antibody, abcam 33985) at 4°C overnight. After being washed 3 times with PBS, the cells were incubated with Cy3B-conjugated donkey anti-rabbit secondary antibody and Alexa 647-conjugated donkey anti-mouse secondary antibody for 2 hours at room temperature, protected from light. The cells were then washed again 3 times and stored in PBS before imaging. Immediately before imaging, the buffer was switched to the STORM imaging buffer containing 10% w/v glucose (Sigma-Aldrich), 0.56 mg/mL glucose oxidase (Sigma-Aldrich), 0.17 mg/mL catalase (Sigma-Aldrich), 0.14M β-mercaptoethanol (Sigma-Aldrich).

### 4. Super-resolution imaging setup

Our experiments were performed on our home-built super-resolution localization microscopy system. It is built upon an Olympus IX71 inverted microscope equipped with four laser lines including 405 nm (DL405-050, CrystaLaser), 488 nm (DL488-150, CrystaLaser), 560 nm (VFL-P-200-560-OEM1, MPB Communications) and 642nm (VFL-P-1000-642-OEM3, MPB Communications). Their intensity was controlled by neutral density filters (NDC-50C-4-A, Thorlabs) and high-speed shutters (LS6S2Z0, Vincent Associates). For super-resolution imaging of tissue section, 642 nm laser with a laser density of 3kW/cm^2^ was used for excitation. The four laser beams were expanded by a 10X beam expander (T81-10X, Newport) and combined by the dichroic mirrors and then focused onto the rear pupil of an oil immersion objective (UPLSAPO 100XO, NA = 1.4, Olympus) by an achromatic lens. A highly oblique-angle illumination was used to suppress the background signal. The emitted fluorescence was collected by the objective, passing through the dichroic mirror (ZT488/640rpc-UF1, Chroma) and a band-pass emission filter (ZET488/640m, Chroma), and then focused by the tube lens and a 0.5X C-mount adapter onto a sCMOS camera (pco.edge 4.2, PCO-Tech), corresponding to a pixel size of 130 nm on the sample plane. A closed-loop piezo nanopositioner (Nano-F100S, Mad City Labs) was used for drift real-time correction tracking the 3D positions of fluorescence nanospheres (F8803, Thermo Fisher Scientific) ^14^. Data acquisition, laser intensity control and drift correction were all integrated in our custom-designed software written in LabVIEW (National Instruments) and MATLAB (MathWorks). We acquired 20,000 frames with an exposure time of 20 ms to ensure the collection of sufficient blinking events. For two-color dSTORM imaging of MCF10A cells, the dichroic mirror was replaced by (ZT405/488/561/640rpc, Chroma) and the emission filter is replaced by (ZET405/488/561/640mv2, Chroma). We acquired 10,000 frames with 642 nm laser with a laser density of 3kW/cm^2^ at an exposure time of 20 ms to ensure the collection of sufficient blinking events.

## Author contributions

H.M. and Y.L. conceived the project. H.M. developed the EVER algorithm, performed the simulation, analysis and cell imaging experiment, and wrote the manuscript. W.J. performed the tissue imaging experiment. J.X. prepared the cell sample. Y.L. supervised the project and wrote the manuscript.

## Acknowledgements

This work is supported by National Institute of Health R01EB016657 and R01CA185363.

